# Piezo1 is required for outflow tract and aortic valve development

**DOI:** 10.1101/528588

**Authors:** Adèle Faucherre, Hamid Moha ou Maati, Nathalie Nasr, Amélie Pinard, Alexis Theron, Gaëlle Odelin, Jean-Pierre Desvignes, David Salgado, Gwenaëlle Collod-Béroud, Jean-François Avierinos, Guillaume Lebon, Stéphane Zaffran, Chris Jopling

## Abstract

**Aims-:** During embryogenesis, the onset of circulatory blood flow generates a variety of hemodynamic forces which reciprocally induce changes in cardiovascular development and performance. It has been known for some time that these forces can be detected by as yet unknown mechanosensory systems which in turn promote cardiogenic events such as outflow tract and aortic valve development. PIEZO1 is a mechanosensitive ion channel present in endothelial cells where it serves to detect hemodynamic forces making it an ideal candidate to play a role during cardiac development. We sought to determine whether PIEZO1 is required for outflow tract and aortic valve development.

**Methods and results-:** By analysing heart development in zebrafish we have determined that *piezo1* is expressed in the developing outflow tract where it serves to detect hemodynamic forces. In particular, we have found that mechanical forces generated during the cardiac cycle activate Piezo1 which triggers nitric oxide to be released in the outflow tract. Consequently, disrupting Piezo1 signalling leads to defective outflow tract and aortic valve development and indicates this gene may be involved in the etiology of congenital heart diseases. Based on these findings, we analysed genomic data generated from a cohort of bicuspid aortic valve patients and identified 3 probands who each harboured a novel variant in *PIEZO1*. Subsequent *in vitro* and *in vivo* assays indicates that these variants behave as dominant negatives leading to an inhibition of normal PIEZO1 mechanosensory activity and defective aortic valve development.

**Conclusion-:** These data indicate that the mechanosensitive ion channel *piezo1* is required for OFT and aortic valve development and, furthermore, dominant negative variants of *PIEZO1* appear to be associated with BAV in humans.

## Introduction

In many species, the onset of circulation precedes the role it will play later in life as an oxygen and nutrient delivery system^1^. As the primitive heart initiates circulation, the forces generated by the blood flow are detected by mechanosensory systems present in the endothelium which lines both the heart and vasculature^2^. These extracellular forces are subsequently converted into intracellular signals which can trigger a variety of cellular responses^3^. Many lines of evidence suggest that the hemodynamic forces generated by the circulatory system act as epigenetic cues to drive developmental processes such as cardiogenesis and valvulogenesis forward^4, 5^. Although a number of mechanosensory systems with the potential to sense circulatory hemodynamics have been identified, our knowledge of how the endothelium detects mechanical stimuli is far from complete^2, 6^. PIEZO1 is a mechanosensitive ion channel present in the cell membrane. When the cell membrane is stretched, this opens the channel and allows an influx of cations^7^. Recently, PIEZO1 has been shown to confer mechanosensitivity to endothelial cells allowing them to detect hemodynamic shear stress and subsequently align themselves in the correct orientation during vasculogenesis^8,9^. Furthermore, deleterious mutations in *PIEZO1* are associated with lymphedema in humans which is caused by defective lymphatic valve development^10^.

The OFT is a transient structure which is extensively remodeled during development and will give rise to a variety of cardiovascular structures including the aortic and pulmonary valves. Although it is well established that hemodynamic blood flow plays a role in OFT development and the formation of the aortic valves^11^, the mechanosensors that detect these forces have remained elusive. However, because of its role in sensing shear stress in the vasculature, *PIEZO1* makes a promising candidate for detecting similar forces in the OFT.

Here we report that disrupting Piezo1 signalling in zebrafish leads to defective development of the OFT and aortic valves. Based on these findings we have been able to identify 3 independent predicted pathogenic *PIEZO1* variants in patients with bicuspid aortic valve disease (BAV). Furthermore, *in vitro* electrophysiological analysis indicates that all variants are dominant negatives which significantly inhibit wild type PIEZO1 activity after stimulation.

## Methods

### Zebrafish strains and husbandry

Zebrafish were maintained under standardized conditions and experiments were conducted in accordance with local approval (APAFIS#4054-2016021116464098 v5) and the European Communities council directive 2010/63/EU. Embryos were staged as described ^9^. The *Tg(fli1a:GFP)y1Tg* was provided by the CMR[B] *Centro de Medicina Regenerativa de Barcelona. The double transgenic line Tg(fli1a:GFP)y1;Tg(cmlc2a:RFP*) was generated in house. All larvae were euthanised by administration of excess anaesthetic (Tricaine).

### Aortic valve imaging

One day prior imaging, larvae were incubated in 0.2 μM BODIPY-FL Ceramide (Invitrogen D3521) in Embryo medium + PTU (0.003% 1-phenyl-2-thiourea). Larvae were then anesthetized with Tricaine (0.16g/L) and mounted in low melting agarose. Imaging was performed with a Zeiss LSM710 two-photon microscope coupled to a Ti:sapphire laser (Spectra-Physics, Santa Clara, CA, USA) and a water immersion 25× objective.

### Electrophysiology

All electrophysiological experiments were performed after 2-6 days of culture for transfected HEK-293T cells seeded at a density of 20 000 cells/35mm dish and after 2-8 hours of culture of freshly dissociated embryonic zebrafish endothelial cells. Dishes were placed on an inverted microscope (DIAPHOT 300, Nikon). Recordings were performed in inside out or cell attached configuration for the patch clamp technique. PIEZO1 currents were elicited by a negative pressure step from 0 to −80mmHg with −10mmHg step increments at −80mV potential. Stimulation protocols and data acquisition were carried out using a PC (Hewlett Packard) with commercial software and hardware (pClamp 10.4) (supplemental information).

### Exome sequencing

The exonic sequences were captured with the Agilent Sure Select All Exon v4 kit (Agilent, Santa Clara, CA, USA) and sequencing was performed on an Illumina HiSeq2000 sequencing apparatus (Illumina, San Diego, CA, USA). Raw Exome sequencing data were aligned against the human reference genome hs37decoy5 and duplicated reads were identified using the SNAP alignment tool version 1.0beta23. For detailed information regarding coverage etc, see (Suppl.table.2). GATK 3.4 was used to perform local indel realignment, score base recalibration and variant calling with the Haplotype Caller. Variations were then selected based on quality criteria using the Variant Filtration module from GATK. Variant annotation (ANNOVAR) and prioritization was performed with the VarAFT software (http://varaft.eu). Prioritization of the filtered-in variants was based on expression in aortic valve according to RNA-seq expression data. Patient recruitment was approved by the Comité de Protection des Patients (13.061).

## Results

### Identifying the zebrafish endothelial *PIEZO1* ortholog

Previous research has identified a zebrafish *PIEZO1* ortholog that does not appear to be expressed in endothelial cells^12^. We have subsequently identified a second *PIEZO1* ortholog, *piezo1b* (Pz1b). At 24 hours post fertilization (hpf), we could detect a weak Pz1b signal in the developing heart tube (Suppl.fig.1.A). By 48hpf a strong expression of Pz1b appeared in the AV canal and OFT (Suppl.fig.1.B). By 4 days post fertilization (dpf) we were able to observe a strong expression of Pz1b in the OFT and vasculature (Fig.1.A,B and Suppl.fig.1.C). Furthermore, we were also able to co-localise Pz1b with GFP labelled endothelial cells in the OFT (Fig.1.C-E). These results indicate that Pz1b is expressed in endothelial cells during zebrafish development, similar to its mammalian counterpart^9^.

**Figure 1.**
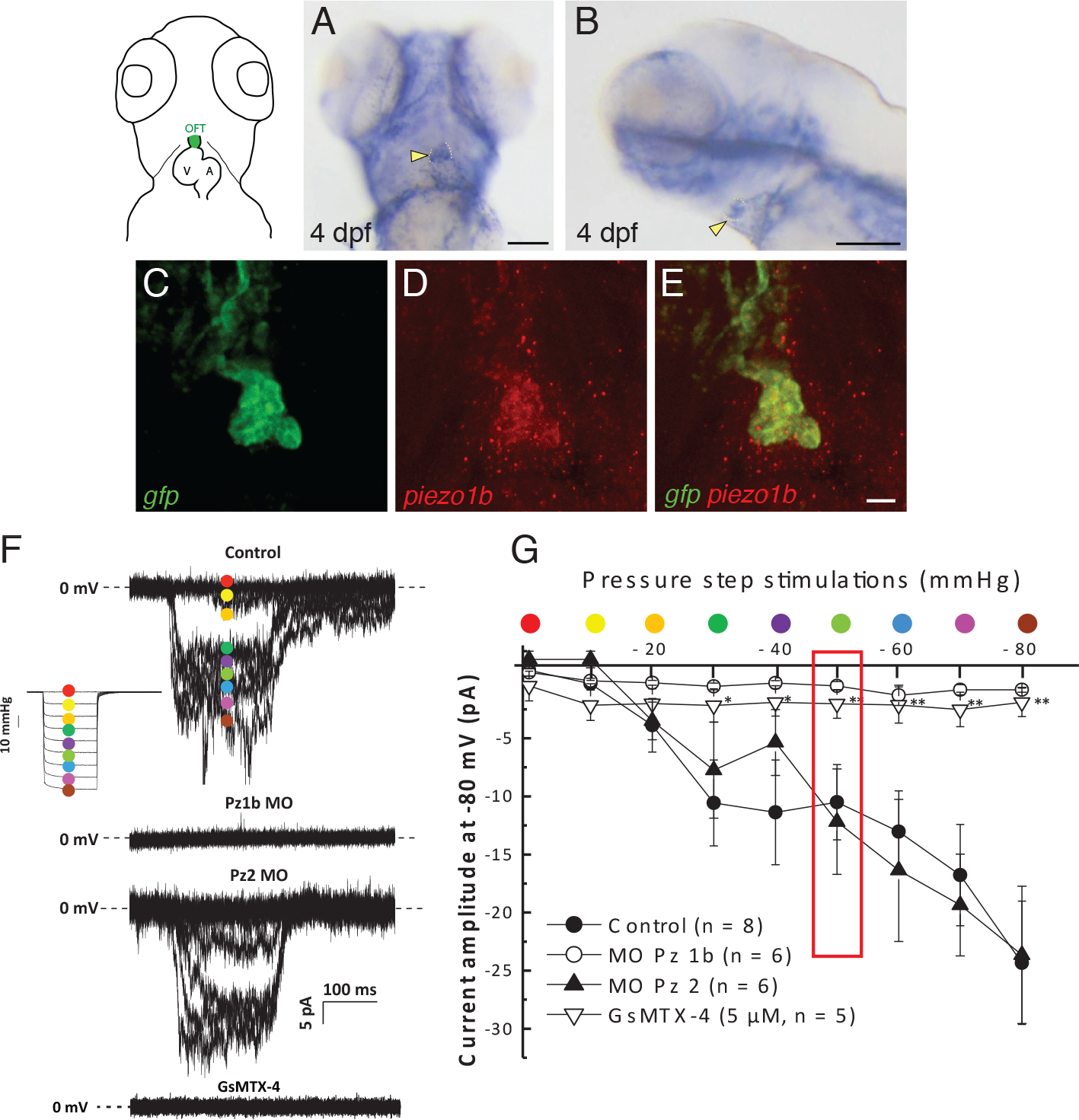
Identifying the zebrafish endothelial *PIEZO1* ortholog. (**A,B**) ISH using an antisense Pz1b probe. Pz1b expression can be detected in the OFT of zebrafish larvae (yellow arrowhead) at 4dpf. (**C-E**) Double fluorescent ISH using antisense GFP (**C**) and antisense Pz1b (**D**) probes, merged image (**E**) indicates endothelial GFP and Pz1b co-localise in the OFT endothelium at 4dpf in *Tg(fli1a:GFP)y1* embryos. (**F,G**) Electrophysiological recordings of mechanosensitive currents from embryonic endothelial cells. Currents were recorded via patch clamp using and inside out configuration by application of negative pressures from 0 to −80mmHg with −10mmHg step increments at −80mV potential. (**F**) Typical current traces recorded from cells isolated from control, Pz1b MO, Pz2 MO injected embryos and cell treated with GsMTx4 *via* the patch clamp pipette. The pressure stimulation protocol is shown in inset. (**G**) Corresponding pressure/current curves showing the activation of the current from control, Pz1b, Pz2 MO cells and GsMTx4 treated cells (* p<0,05, ** p<0.01, student’s t-test). The red rectangle shows the value of the current amplitude for the three test conditions at −50 mmHg step stimulation. Scale bars: (**A,B**) 100μm (**E**) 10μm.

We next sought to determine whether Pz1b is a functional mechanosensitive ion channel in the developing zebrafish endothelium. To achieve this, we performed electrophysiological analysis of cultured zebrafish endothelial cells subjected to mechanical stimulation (Fig.1.F,G). Furthermore, we found that knockdown of Pz1b abolishes mechanically induced currents which was not the case when we targeted *piezo2*^13^ (Fig.1.F,G). As an additional control, we performed the same electrophysiological analysis on WT endothelial cells treated with the PIEZO channel inhibitory peptide GsMTx4^14^ and also observed a reduction in the amplitude of mechanically induced currents (Fig.1.F,G). These data indicate that Pz1b is a functional mechanosensitive ion channel present in the endothelial cells of developing zebrafish embryos.

### *Piezo1b* is involved in cardiovascular development

Knockdown of Pz1b produced a phenotype characterized by defective cardiogenesis and associated oedema at 72hpf (Suppl.Fig.2. A-D’ and Suppl.fig.3.G-H’). Although we ensured the specificity of the observed phenotype to the knockdown of Pz1b by employing two different morpholinos targeting the same gene, we also analysed mRNA splicing, morpholino synergy and generated a CRISPR/Cas9 Pz1b knockout (Suppl.information and suppl.figs.S3 and S4). We next sought to rescue the Pz1b morphant phenotype using mouse *piezo1* mRNA (mPz1) which is not targeted by the Pz1b-MO1. In this manner, we found that co-injection of Pz1b-MO1 with 20pg of mPz1 RNA could rescue the cardiac defects observed in Pz1b morphants (Suppl.fig.3.C-F). Analysis of a number of physiological parameters indicates that Pz1b morphants exhibit a decreased cardiac output along with blood regurgitation through the AV canal (Suppl.fig.2.E-I and suppl.information and suppl.fig.S6 and movies S1,2). Taken together these data indicate that loss of Pz1b results in defective cardiac development.

**Figure 2.**
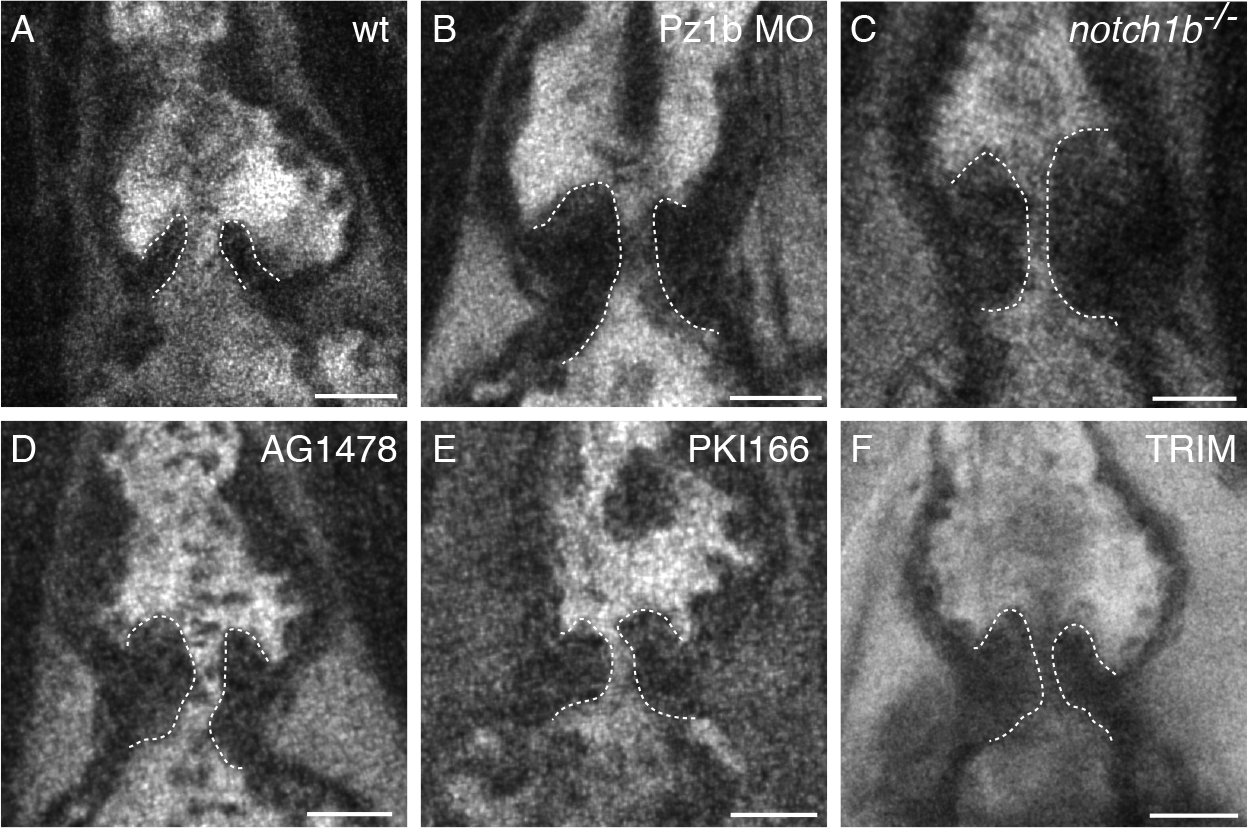
*Piezo1b* is required for aortic valve development. 2 photon images of aortic valves in 7dpf zebrafish larvae labelled with BODIPY. **(A)** Representative image of the aortic valves in a WT larvae, valves are outlined with a dashed white line (n=8). **(B)** A Pz1b morphant (n=6). **(C)** A *notch1b* mutant (n=11). **(D)** A AG1478 treated larvae (n=6). **(E)** A PKI166 treated larvae (n=6). **(F)** A TRIM treated larvae (n=7).

### *Piezo1b* is required for OFT development

To determine whether OFT development has been affected, we analysed this structure in WT and Pz1b morphants. At 48hpf, we could not detect any discernible differences between WT and Pz1b morphant embryos (Suppl.fig.2.J,K). However, by 72hpf, the WT OFT had developed into a “pear shaped” structure which was noticeably wider at the interface with the myocardium when compared to the Pz1b morphant OFT (Suppl.fig.2.L-P). Furthermore we were able to rescue this defect by co-injecting mouse *piezo1* RNA (suppl.fig.S3.Q,R).

Because the OFT is directly adjacent to the ventricle, it is most likely subjected to some of the most extreme hemodynamic forces which in turn could be detected by Pz1b to subsequently initiate the further development of this structure. To test this premise we analysed the dynamics of OFT in zebrafish larvae using confocal microscopy. At 48hpf we were able to observe the OFT stretch and relax during the cardiac cycle. At the end of systole, the OFT achieved a maximum peak inside diameter of 16.87μm (+/−0.24 SEM) (Suppl.fig.2.Q-S and movie S3). We also made similar measurements of the dorsal aorta as a comparison and found that its diameter increased by only 0.72μm (+/−0.20 SEM) during systole (Fig.5.S and movie S4). This indicates that indeed hemodynamic forces produce a strong dynamic response in the OFT endothelium in comparison to other regions of the vasculature.

### *Piezo1b* is required for aortic valve development

During cardiac development, the aortic valves will form in the OFT, a process reliant on hemodynamic forces. Based on this notion we analysed the developing aortic valves to determine whether Pz1b is involved in this process. To achieve this we labelled 7dpf larvae with BODIPY and imaged them using 2 photon microscopy, a technique which has previously been employed to analyse the development of the atrioventricular valves in zebrafish^15^. Our analysis of WT larvae indicates that at this time point two defined leaflets have formed which function effectively to regulate the flow of blood as it is ejected from the ventricle (Fig.2A and Movie.S5). In comparison, the valves in Pz1b morphants are highly dysmorphic and appear to be enlarged and misshapen (Fig.2.B and Movie.S6). Previous research in mammals and humans has determined that *NOTCH1* is required for aortic valve development and mutations in this gene are associated with BAV^16^. In this regard we next analysed the aortic valves in *notch1b* KO zebrafish larvae. Similar to Pz1b morphants, the aortic valves in *notch1b* mutants are also highly dysmorphic compared to WT controls (Fig.2.C and Movie.S7). The *epidermal growth factor receptor* (EGFR) gene has also been implicated in the etiology of BAV^17^, furthermore previous research in zebrafish has indicated that chemical inhibition of EGFR using either PKI166 or AG1478 results in defective OFT development^18^. Based on this we treated zebrafish embryos with either inhibitor for 7 days and again assessed aortic valve development. As with Pz1b morphants and *notch1b* mutants, inhibition of EGFR results in dysmorphic aortic valves (Fig.2.D,E). Lastly, *Endothelial Nitric Oxide Synthase* (eNOS) has also been linked to BAV in mice^17^, while treatment of zebrafish embryos with the NOS inhibitor 2-trifluoromethylphenyl imidazole (TRIM) results in defective cardiac development^19^. Therefore, we also treated zebrafish larvae with TRIM and analysed their aortic valves as before and found that TRIM treated larvae also display defective aortic valves (Fig.2.F). Taken together these data indicate that loss of Pz1b results in defective aortic valve development and, furthermore, the resulting phenotype is reminiscent of the valve phenotype caused by disrupting BAV associated genes/proteins in zebrafish.

### *Piezo1b* regulates nitric oxide production and extracellular matrix composition in the outflow tract

During embryonic zebrafish development, there is a pronounced and sustained release of nitric oxide (NO) in the OFT, coincident with increasing hemodynamic load^20, 21^. To determine whether Pz1b is involved in NO release, we utilized a DAF-FM DA assay to detect the NO signal produced in the OFT ^20, 21^. Analysis of control 72hpf embryos indicates a clear NO signal in the developing OFT (Suppl.fig.5.A-B’and 5.I). Conversely, the NO signal was noticeably absent in the OFT of Pz1b morphants (Suppl.fig.5.C-C’and 5.I). We next assessed whether ablation of erythrocytes with phenylhydrazine (PHZ) to reduce shear stress, or reducing the heart rate substantially (108.3bpm +/−2.1SEM, n=10) by treatment with 10mM 2,3-butanedione monoxime (BDM) ^22, 23^, also had the same effect on NO production in the OFT (Suppl.fig.5.D-E’ and 5.I). Although each of these conditions leads to a significant reduction in NO signal, only complete cessation of the heart beat using 15mM BDM abolishes the NO signal (Suppl.fig.5.F-F’ and 5.I). To further confirm these results we also analysed NO production in *troponin t2* (*tnnt2*) morphants which lack any discernible heartbeat^24^. Similarly to arresting the heart using BDM treatment, we also found that the loss of heart beat observed in *tnnt2* morphants also significantly reduces the production of NO in the OFT (Suppl.fig.5.G-G’ and 5.I). Lastly, we also treated WT 72hpf embryos with the PIEZO1 inhibitor, GsMTx4, for 4 hours and found that this significantly reduced the amount of NO produced in the OFT (Suppl.fig.5.H-H’ and 5.I). This data indicates that hemodynamics forces produced in the OFT induce NO production and that Pz1b plays a role in detecting these forces. Previous research has identified that the extracellular matrix (ECM) component, Elastin, is produced in the OFT where it provides the elasticity to cope with increasing hemodynamic load and to dampen the force of blood as it is ejected from the ventricle^25^. We performed immunohistochemistry on 72hpf zebrafish larvae using a previously described ElastinB (ElnB) antibody^25^, and were able to determine that ElnB is produced by cells which ensheathe the developing OFT endothelium (Suppl.fig.5.J-L). Where these cells originate from is at present ambiguous however, both neural crest cells and second heart field progenitors do appear to contribute^26, 27^. We next assessed whether hemodynamic forces could play a role in triggering ElnB production. To achieve this we performed ISH using an *elnB* riboprobe on 72hpf zebrafish larvae. Under normal conditions we could detect a clear and specific expression of *elnB* in the OFT (Suppl.fig.5.M). However, analysis of tnnt2 morphants which lack a heartbeat, and so are devoid of hemodynamic forces, revealed an obvious loss of *elnB* expression in the OFT (Suppl.fig.5.N). Similarly, in Pz1b morphants, *elnB* expression is also appreciably reduced (Suppl.fig.5.O) and lastly, in embryos treated with TRIM to inhibit NO production, *elnB* expression in the OFT is clearly reduced (Suppl.fig.5.P). These data indicate that *elnB* expression in the OFT is hemodynamically dependent and that both Pz1b and NO are required for this process. Previously we have shown that the ECM component AggrecanA (Acana) is also expressed in the same population of cells as ElnB and in a hemodynamically dependent manner^28^. Furthermore, are data indicates that reduced *AGGRECAN* expression is associated with type 0 BAV in humans. Due to this, we next analysed whether Pz1b could regulate the expression of *acana*. ISH using an *acana* riboprobe on 72hpf WT zebrafish larvae indicates that *acana* is strongly expressed in the OFT at this timepoint (Suppl.fig.5.Q). Conversely, when Pz1b is knocked down the expression of *acana* in the OFT is clearly reduced (Suppl.fig.5.R). Taken together our data highlights a potentially novel signaling pathway initiated by hemodynamic forces in the OFT which are subseqeuntly detected by Pz1b, this, in turn, triggers NO to be released, which subsequently initiates the production of ElnB and AcanA (Suppl.fig.7).

### Identification of human variants in *PIEZO1* associated with BAV

To determine whether mutations in *PIEZO1* are associated with valvulopathies such as BAV, we examined whole exome sequence data generated in-house from 19 patients diagnosed with isolated BAV. In parallel, we also analysed whole exome sequence data from 30 BAV patients provided by the National Heart, Lung, and Blood Institute (NHLBI) Bench to Bassinet Program: The Pediatric Cardiac Genetics Consortium (PCGC) dataset (dbGaP accession phs000571.v3.p2.). In this manner, we were able to identify 3 independent nonsynonymous variants in *PIEZO1*: p.Tyr2022His (c.6064T>C; p.Y2022H; located in a transmembrane C-terminal domain (in-house analysis)), p.Lys2502Arg (c.7505A>G; p.K2502R; located in the cytoplasmic C-terminal region of PIEZO1 (in-house analysis)) and p.Ser217Leu (c. 650C>T; p.S217L; located in N-terminal region (PCGC analysis)) (Fig.3.A-C). The heterozygous p.Y2022H and p.K2502R variants were subsequently validated by Sanger sequencing (Fig.3.B,C). Although this was not possible for the p.S217L variant, we were able to determine that one of the parents also harboured this mutation indicating that the proband is heterozygous for this variant. According to gnomAD^29^, all 3 variants are considered rare, with minor allele frequencies <1% (Table.1). By comparing PIEZO1 orthologs in different species it is apparent that Tyr2022, Lys2502 and Ser217 are evolutionary conserved amino acids (Fig.3.F). To determine the potential functional consequences these variants have on PIEZO1, they were analysed using the CADD^30^, Mutation Taster2^31^ and UMD Predictor programs^32^ (Table.1). In this manner, all 3 variants are predicted to be pathogenic. Pedigree analysis for all 3 probands can be found in the supplemental information.

**Figure 3.**
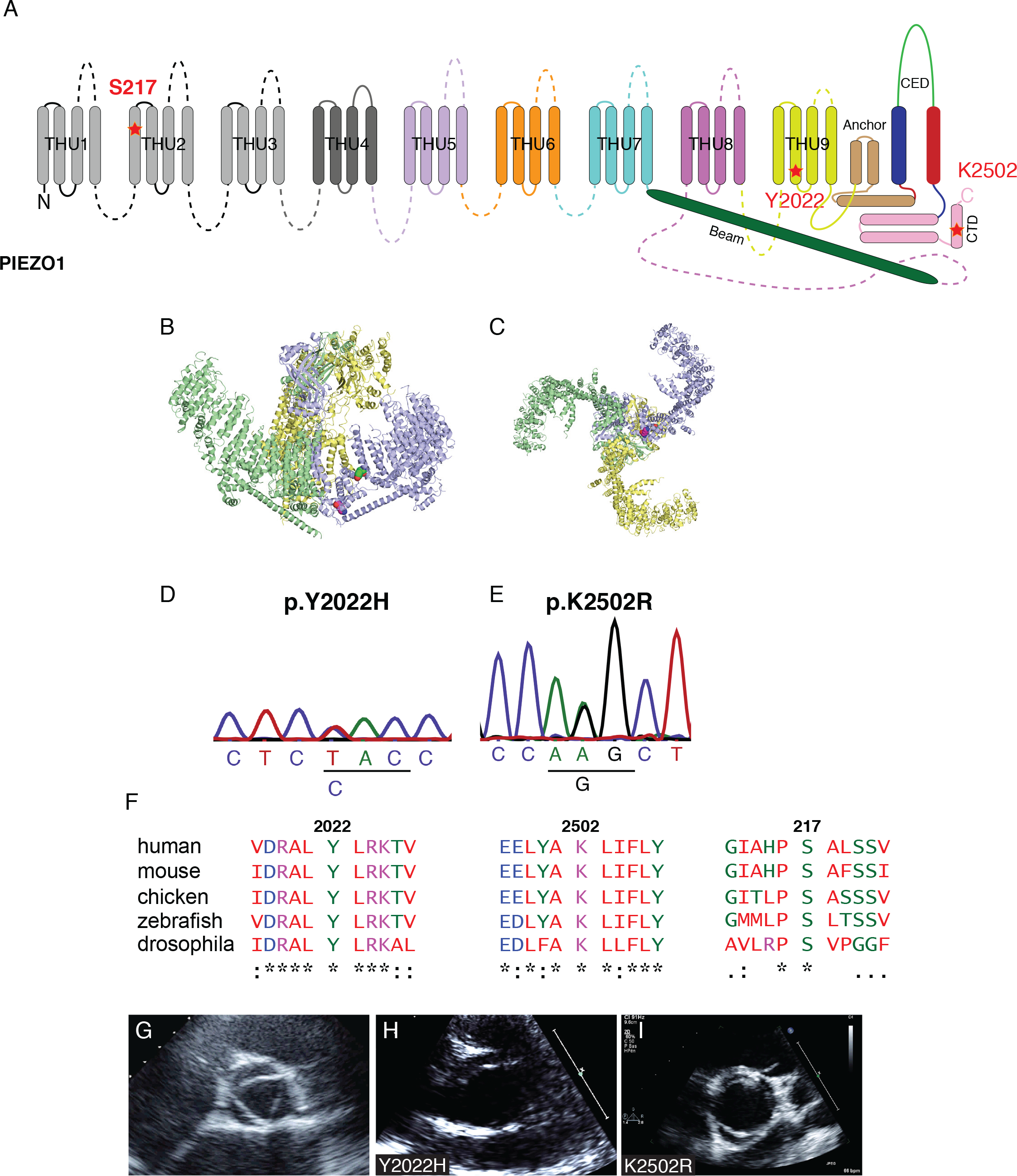
Identification of variants in *PIEZO1* associated with BAV. (**A**) Schematic representation of human PIEZO1 protein, the position of each variant is indicated by a red dot (adapted from *Zhao et al^49^*). (**B,C**) Cartoon representation of the mouse mechanosensitive Piezo1 channel 3D Model (PDB 5Z10). The corresponding residues K2502 (Magenta) and Y2022 (Green) identified in this study as subject to genetic variability are represented as spheres. (**D,E**) Sequence chromatographs showing heterozygous variants c.6064T>C (p.Y2022H; B) and c.7505A>G (p.K2502R; C) in genomic DNA taken from the affected BAV patients. (**F**)Protein alignments for all 3 variants. Note the strong evolutionary conservation for Tyr2022, Lys2502 and Ser217 residues. (**G-I**) Representative echocardiogram of the aortic valve from an unaffected individual (**G**) and the BAV patients carrying out the c.6064T>C; p.Y2022H variant (**H**) and c.7505A>G; p.K2502R variant (**I**). Note the 3 valve leaflets in the unaffected individual (**G**) compared to the 2 leaflets in the BAV patients (**H** and **G**).

**Table 1.** Prediction analysis of *PIEZO1* mutations.

| *Piezo1* mutation | CADD-PHRED | Mutation Taster | UMD Predictor | gnomAD allele frequencies |
| --- | --- | --- | --- | --- |
| p.Y2022H<br>(c.6064T>C) | Damaging (27.1) | Damaging (0.58) | Pathogenic (99) | 3.336 10<sup>-5</sup> |
| p.K2502R<br>(c.7505A>G) | Damaging (25.6) | Damaging (0.80) | Probably pathogenic (69) | 6.498 10<sup>-3</sup> |
| p.S217L<br>(c.650C>T) | Damaging (25.6) | Damaging (0.54) | Pathogenic (78) | Not reported |

### Functional analysis of BAV associated variants in *PIEZO1*

To determine whether the identified variants affected PIEZO1 protein function, we performed mechano-electrophysiology analysis of HEK-293T cells transfected with either wildtype (WT) human *PIEZO1* or the p.Y2022H, p.K2502R and p.S217L variants. In this manner we were able to detect changes in current from HEK cells transfected with wildtype human *PIEZO1* (Fig.4.A,E). Conversely, cells which had been transfected with either p.Y2022H, p.K2502R or p.S217L *PIEZO1* variants showed significantly reduced mechano-stimulated currents (Fig.4.B-E). One explanation for the reduction in *PIEZO1* activity associated with these variants is that the mutations result in a reduction of cell surface expression. To assess this possibility we performed immunohistochemistry (IHC) on HEK-293T cells transfected with either wildtype human *PIEZO1* the p.Y2022H, p.K2502R or p.S217L *PIEZO1* variants. Confocal images analysis of transfected cells indicates that none of the variants appear to significantly affect the localization of PIEZO1 to the cell surface when compared to wildtype PIEZO1 (Fig.4.F). To confirm this observation we measured the fluorescent intensity across the cell membrane. In this manner, wildtype PIEZO1 and all of the variants show a clear peak of intensity at the cell surface which drops sharply on the extracellular and intracellular sides when compared to GFP, which is expressed throughout the cell (Fig.4.G). This shows that aberrant trafficking of PIEZO1 to the cell surface is not the reason for the observed reduction in activity. These data indicate that the variants p.Y2022H, p.K2502R and p.S217L have deleterious effects on PIEZO1 protein function.

**Figure 4.**
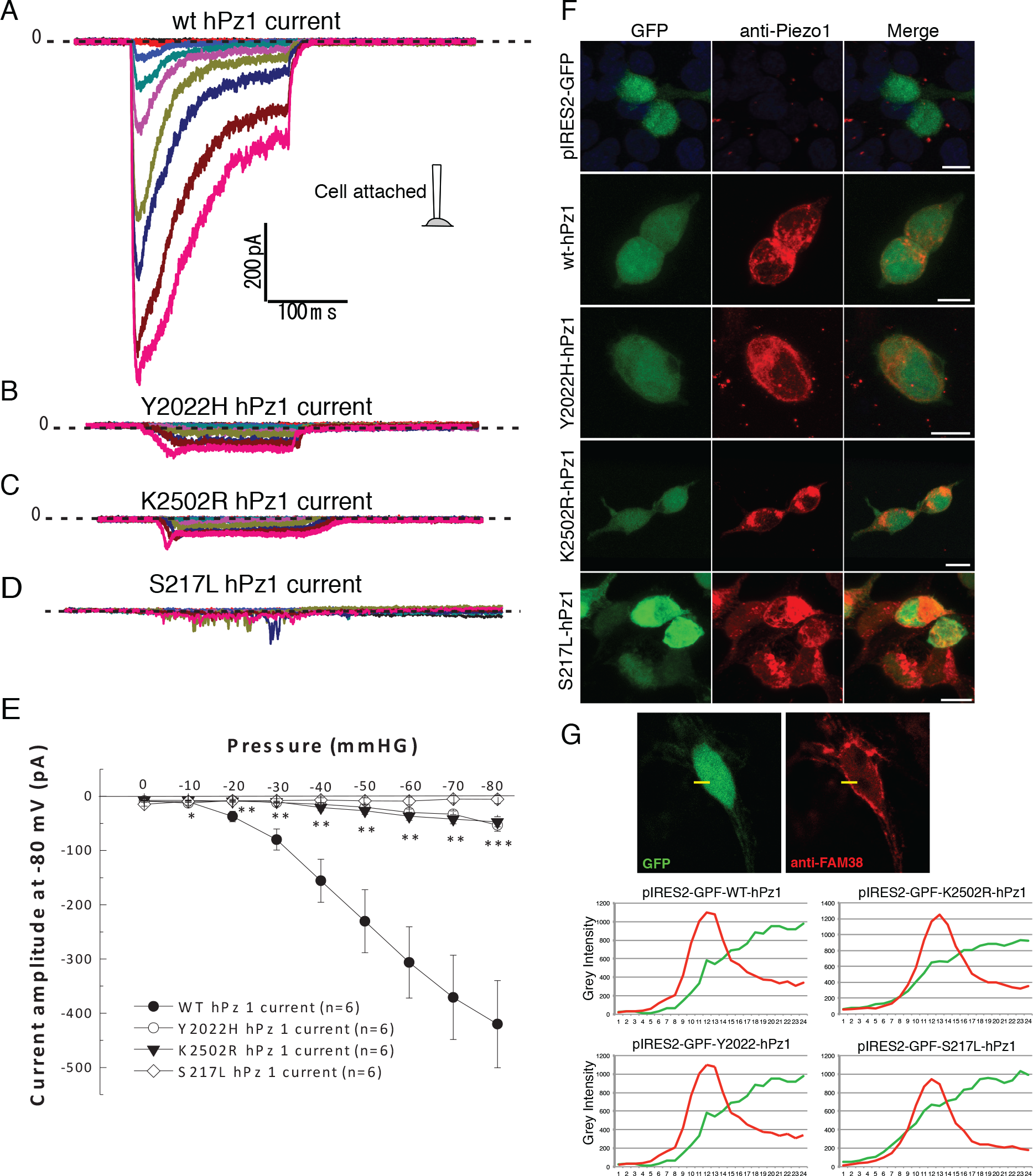
Functional analysis of BAV associated variants in *PIEZO1*. Electrophysiological recordings of mechanosensitive currents from all 3 *PIEZO1* variants associated with BAV. Typical current traces recorded from HEK cells transfected with either wildtype (WT) human *PIEZO1* (hPz1) (**A**), human *PIEZO1* c.6064T>C (p.Y2022H) variant (**B**), human *PIEZO1* c.7075A>G (p.K2502R) (**C**) variant or human *PIEZO1* c.650C>T (p.S217L) variant (**D**). Currents were recorded *via* patch clamp using a cell attached configuration by application of negative pressures from 0 to −80mmHg with −10mmHg step increments at −80mV potential. (**E**) Corresponding pressure/current curves showing the activation of the current from wildtype hPz1, p.Y2022H, p.K2502R and p.S217L (* p<0,05, ** p<0.01, *** p<0.001 student’s t-test). (**F**) Immunohistochemistry using an anti PIEZO1 antibody on HEK 293T cells transfected with either empty vector (pIRES2-GFP), wildtype human *PIEZO1* (wt-hPz1), human *PIEZO1* c.6064T>C (p.Y2022H) variant (Y2022H-hPz1), human *PIEZO1* c.7075A>G (p.K2502R) variant (K2502R-hPz1) or human *PIEZO1* c.650C>T (p.S217L) variant (S217L-hPz1). **(G)** Cell surface immunofluorescent analysis. Fluorescent intensity was measured in ImageJ by drawing a line through the cell membrane (yellow bar). Graphs plotting the fluorescent intensity at the cell surface for WT and variant *PIEZO1*. In all cases there is a clear peak of RFP fluorescent intensity which tapers off on either side of the cell membrane. The control GFP signal is continuous. Scale bars (F) 10μm.

**Figure 5.**
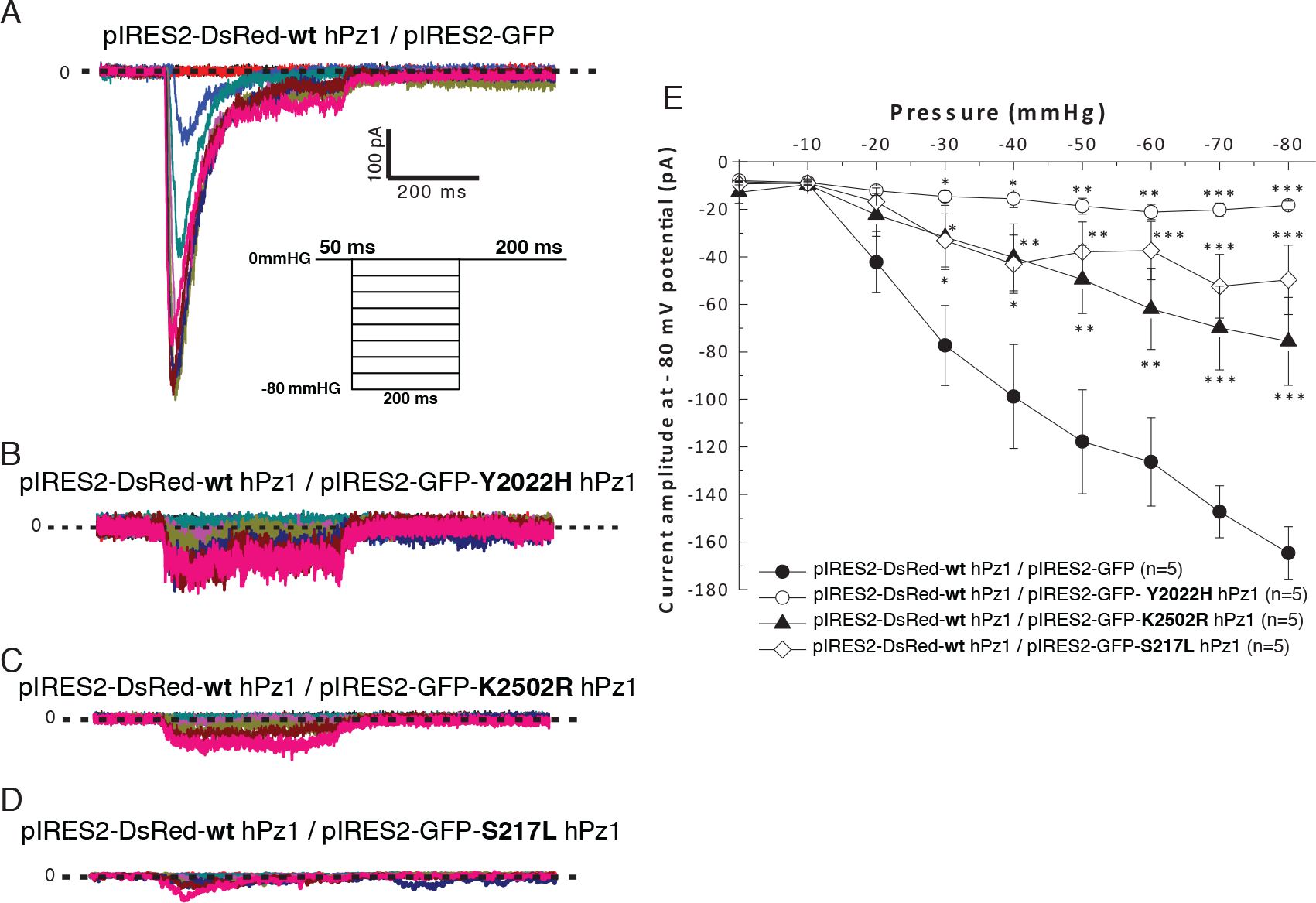
BAV associated *PIEZO1* variants are dominant negatives. Electrophysiological recordings of mechanosensitive currents from co-transfection of 3 *PIEZO1* variants with wildtype human *PIEZO1*. Typical current traces recorded from HEK cells co-transfected with either wildtype human *PIEZO1* (hPz1) and empty pIRES2-GFP (**A**), wildtype human *PIEZO1* (hPz1) and human *PIEZO1* c.6064T>C (p.Y2022H) variant (**B**), wildtype human *PIEZO1* (hPz1) and human *PIEZO1* c.7075A>G (p.K2502R) variant (**C**) or wildtype human *PIEZO1* (hPz1) and human *PIEZO1* c.650C>T (p.S217L) (**D**). Currents were recorded *via* patch clamp using a cell attached configuration by application of negative pressures from 0 to −80mmHg with −10mmHg step increments at −80mV potential. (**E**) Corresponding pressure/current curves showing the activation of the current from wildtype hPz1/empty pIRES2-GFP, wildtype hPz1/p.Y2022H, wildtype hPz1/p.K2502R and wildtype hPz1/p.S217L (* p<0,05, ** p<0.01, *** p<0.001 student’s t-test).

### p.Y2022H, p.K2502R and p.S217L *PIEZO1* variants are dominant negative isoforms

Because all 3 probands are heterozygous for their respective variants, we next sought to ascertain whether the association with BAV was due to haploinsufficiency or to a possible dominant negative effect of these mutations. When wildtype human *PIEZO1* was co-transfected with either p.Y2022H, p.K2502R or p.S217L *PIEZO1* variants, there was a significant decrease in current amplitude compared to the control (Fig.5.A-E). Taken together these data indicate that the p.Y2022H, p.K2502R and p.S217L *PIEZO1* variants act as dominant negatives.

Next we sought to determine whether forced expression of the dominant negative human *PIEZO1* variants in the zebrafish endothelium could also affect aortic valve development. To achieve this we generated a transgenic construct using the endothelial specific *fliEP* promoter^33^ to drive expression of either WT *PIEZO1* or the variants specifically in endothelial cells *in vivo*. In this manner we were able to determine that expressing any of the dominant negative variants (p.Y2022H, p.K2502R or p.S217L) also had a significant impact on aortic valve development when compared to WT *PIEZO1* expression. To assess this in more detail we performed a quantification of the defective valves by dividing the length of either leaflet by its width (Fig.6.A,B). In this manner we found that all the variants significantly reduced the overall length/width ratio in both leaflets when compared to the WT *PIEZO1* expressing larvae. This indicates that expression of either Y2022H, p.K2502R or p.S217L *PIEZO1* disrupts aortic valve development in zebrafish.

**Figure 6.**
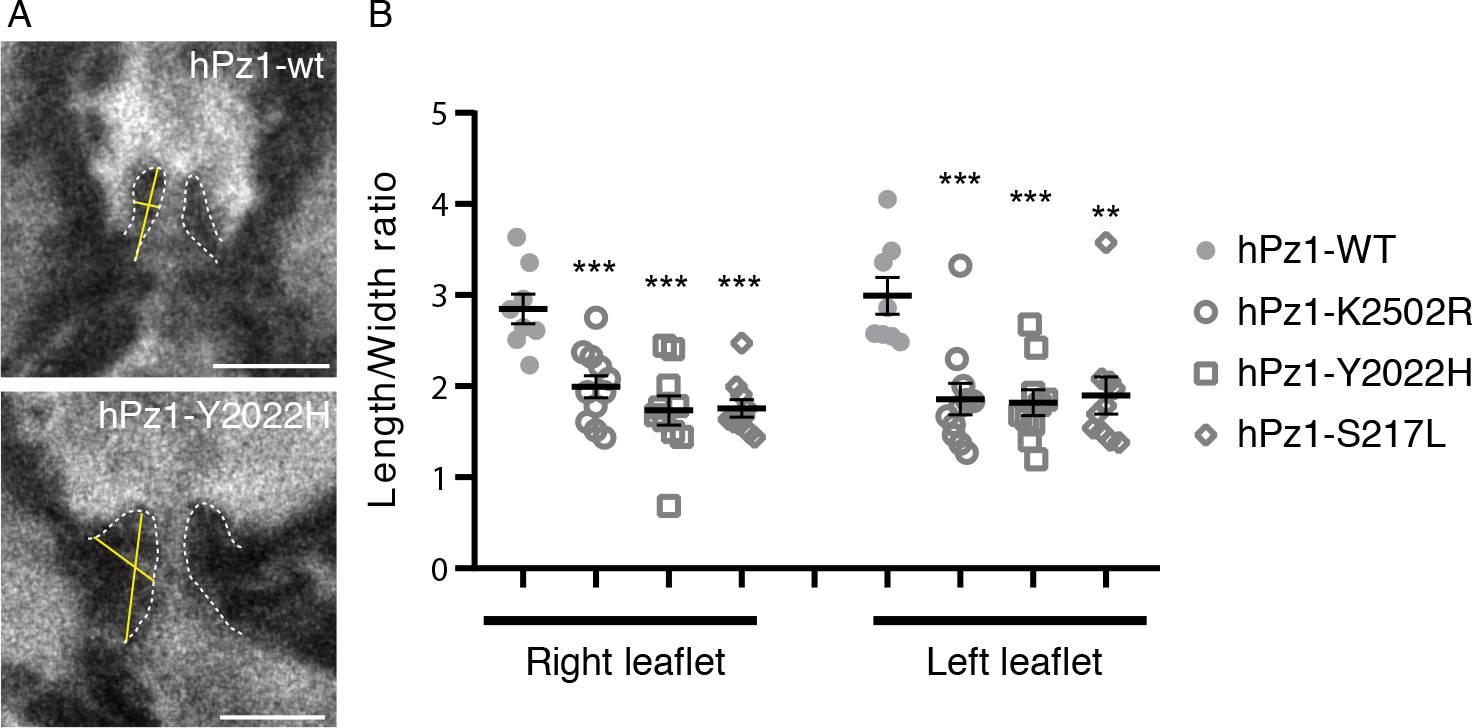
Expression of *PIEZO1* variants *in vivo* disrupts aortic valve development. **(A)** Representative images of aortic valve leaflet quantification in a 7dpf larvae expressing either WT human *PIEZO1* (hPZ1-wt) or the Y2022H variant of human *PIEZO1* (hPZ1-Y2022H), ratios were calculated by measuring the length of the leaflet from the lower edge to the tip, the width was measured at a point halfway along the length (yellow lines). **(B)** Graph depicting the length to width ratio for the left and right valve leaflets measured in 7dpf larvae expressing either WT (n=8), K2502R (n=11), Y2022H(n=10) or S217L (n=10) human *PIEZO1*. Error bars indicate SEM, ANOVA and Dunnet’s multiple comparisons test P<0.01(all samples), student’s unpaired homoscedastic two tailed t-test **P<0.01. Scale bars-20μm.

## Discussion

Despite the differences in cardiac physiology, early OFT development is highly conserved between mammals and zebrafish, in particular this region will give rise to the aortic valves. Recently, it has been established that the coordinated actions of the mechanosensitive ion channels TRPV4 and TRPP2 are required to promote *klf2a* expression in the AV canal and subsequently drive valve morphogenesis in this region^34^. It appears that Pz1b may play a similar mechanosensory role in the OFT where it is required to trigger NO production in response to increasing hemodynamic load. How Pz1b triggers NO production is at present unclear, however previous research has highlighted a feedback mechanism that involves the release of ATP from the endothelium which activates a P2Y2 signalling cascade resulting in NOS activation^35^. Interestingly, treating zebrafish embryos with the Nitric Oxide Synthase (NOS) inhibitor TRIM also disrupts OFT development and leads to a similar cardiac phenotype^36^ while mice deficient in Endothelial Nitric Oxide Synthase (eNOS) develop BAV^37, 38^. Our results are also in line with mammalian data that indicates a direct regulation of eNOS by PIEZO1^8^. *NOTCH1* is one of the few genes directly linked to BAV in humans and evidence suggests that NO can also regulate NOTCH1 signaling during aortic valve development. Compound mutant mice which are *eNOS*^−/−^;*Notch1*^+/−^ show a dramatic increase in the prevalence of BAV when compared to either *eNOS*^−/−^ or *Notch1*^+/−^ alone^39^. Our own data indicates that the zebrafish *NOTCH1* ortholog *notch1b* also regulates aortic valve development. Indeed, zebrafish *notch1b* mutants display dysmorphic valves. Importantly, we also observed a very similar aortic valve phenotype in Pz1b morphant larvae indicating that this gene is also required for aortic valve development. Future investigation will be aimed trying to ascertain whether Pz1b mediated NO release acts synergistically with Notch1b during valve development.

In humans, disruption of ECM components has been linked to a variety of OFT pathologies such as aortic stenosis (AS) and bicuspid aortic valves (BAV) ^40–42^. Mutations and decreased expression of *ELASTIN* have been linked to supravalvar aortic stenosis (SVAS), a condition which leads to the narrowing of the aorta adjacent to the aortic valve ^41^. Furthermore, a chromosomal microdeletion which includes *ELASTIN* causes Williams Beuren syndrome which manifests with a variety of developmental defects including SVAS and BAV^40^. Similarly, *ELASTIN* haploinsufficiency in mice also leads to progressive aortic valve degeneration^43^. In chick embryos, ELASTIN production is initiated at day 3 by cells which surround the endothelium of the aorta directly adjacent to the myocardium, before spreading throughout the vasculature ^44^. In zebrafish, Elastin production in the OFT also commences at around 3 days post fertilisation and coincides with cardiogenic events such as coordinated contraction and AV canal development that place increasing hemodynamic loads on the OFT. Here we provide evidence of a novel signalling cascade initiated by hemodynamic forces in the OFT which are detected by Pz1b and ultimately leads to the expression of ECM components, such as Elastin. Future studies will be required to determine whether BAV patients who harbour deleterious *PIEZO1* variants have a decreased expression of ECM components. It will also be interesting to determine whether other ECM components such as COLLAGEN are also regulated by a similar mechanism.

We have also identified 3 pathogenic PIEZO1 variants associated with BAV in humans. Although the variants we have identified inhibit PIEZO1 function, for the p.Y2022H mutant, the proband’s twin, presenting with a tricuspid valve, carried the same variant. This could be due to incomplete penetrance linked to modifiers. For example, mutations in *NOTCH1* have been linked to CHD, however the BAV phenotype is not fully penetrant^16^. Furthermore, it has recently been shown that in monozygotic twins, despite the absence of any pathogenic genetic differences between them, only one of the pair developed BAV^45^. It should be noted that inhibitory variants in *PIEZO1* are also associated with a novel form of hereditary lymphedema^46, 47^. Although neither report indicates the presence of BAV, it is also unclear whether the patients underwent echocardiography to detect this. However, it is interesting to note that in other conditions in which lymphedema is present, such as Turner syndrome, there is an increased prevalence of BAV (28.4%)^48^. Whether this is also the case for the *PIEZO1* form of lymphedema will require more detailed and expansive analysis of this condition.

## Acknowledgements

We would like to acknowledge Dr Matteo Mangoni and Dr Joel Nargeot for their input and support. We would like to thank Prof Ardem Patapoutian for the kind gift of h*PIEZO1*-pIRES2-GFP. We would like to thank Dr Emmanuel Bourinet for the kind gift of m*Piezo1*. We would also like to acknowledge the Montpellier MGX Genomix platform for their input and support. Part of this data was generated by the Pediatric Cardiac Genomics Consortium (PCGC), under the auspices of the National Heart, Lung, and Blood Institute’s Bench to Bassinet Program <http://www.benchtobassinet.org/>. The Pediatric Cardiac Genomics Consortium (PCGC) program is funded by the National Heart, Lung, and Blood Institute, National Institutes of Health, U.S. Department of Health and Human Services through grants U01HL098123, U01HL098147, U01HL098153, U01HL098162, U01HL098163, and U01HL098188. This manuscript was not prepared in collaboration with investigators of the PCGC, has not been reviewed and/or approved by the PCGC, and does not necessarily reflect the opinions of the PCGC investigators or the NHLBI. We thank Anthony Pinot from the *in vivo* imaging platform IPAM-Biocampus Montpellier.

## Supplemental information

### Pedigree analysis of BAV patients harboring variants in *PIEZO1*

The BAV phenotype found in the p.Y2022H and p.K2502R probands was diagnosed by echocardiography (Fig.3.G-I). The proband carrying the p.Y2022H (c.6064T>C) variant was diagnosed in childhood with several congenital malformations including a BAV without ascending aorta aneurysm, a left arm agenesis, a bicornate uterus and a urogenital malformation. Physical examination revealed an aortic murmur at the third intercostal space. Transthoracic echocardiography showed a BAV (Fig.3.H and movie S8) with a raphe between the right and the non-coronary leaflets (so called I, R/NC, regarding the Sievers classification) and a mild-to-moderate aortic regurgitation (Effective regurgitant orifice area = 16mm^2^) due to a prolapse of the anterior leaflet. Pedigree analysis showed that the proband’s twin carried the same variant. However, ultrasound and examination revealed no congenital malformation in this individual. No other living relatives were positive for the variant or had cardiac complaints.

The proband carrying the p.K2502R (c.7505A>G) variant was incidentally diagnosed at 13 years old. Family medical history revealed that his mother had a mitral valve prolapse. However, we have no information on the mother’ genotype. Physical examination revealed a 3/6 holodiastolic murmur. Transthoracic echocardiography showed a BAV (Fig. 3.I; and movie S9) with a raphe between the right and the left coronary leaflets (I, R/L, regarding the Sievers classification) and a moderate aortic regurgitation (Effective regurgitant orifice area = 20mm^2^) due to a prolapse of the anterior leaflet. There was no dilatation of the ascending aorta. Left ventricular ejection fraction was 70%.

The proband (dbGaP Subject ID-711051) harboring the p.S217L (c. 650C>T) variant was diagnosed in adolescence (age 17.3 years old) by echocardiography with a BAV R/NC fusion. Further cardiac analysis revealed no other associated cardiac defects (cardiac situs, systemic vein, hepatic vein, pulmonary vein, right atrium, atrial septum, left atrium, atrioventricular junction, tricuspid valve, mitral valve, right ventricle, left ventricle, pulmonary valve, coronary arteries, pulmonary arteries and aorta are normal). The proband is a caucasian male born in the United States of America, he stands at 162.2cm tall and weighs 117kg, he presented with no other extra cardiac abnormalities. His mother and father are caucasian Americans and were 22 and 26 years of age respectively at the birth of the proband. The mother also harbors the same p.S217L variant. Although the mother has no reported history of heart disease, we have no information regarding whether or not she has undergone echocardiography to detect BAV (a limitation of the study recognized by the PCGC). No further information regarding the mother is available.

### Morpholinos

The sequences of the injected MOs are the following:

Pz1b MO1: 5′-TCTGTTGCTGGATTCTGTGAATCAT-3’-5ng
Pz1b MO2: 5′-ACCCATGATGCTGCAACACACACAC-3’-4ng
*tnnt2* MO: 5′-CATGTTTGCTCTGATCTGACACGCA3’ -3ng

### DNA microinjection

The fli1ep:h*PIEZO1* constructs were generated using the Tol2 kit^1^. 478 p5Efli1ep was a gift from Nathan Lawson (Addgene plasmid # 31160; http://n2t.net/addgene:31160;RRID:Addgene_31160). Wildtype variant h*PIEZO1* were subcloned from pIRES2-GFP into pME-MCS by restriction digestion and ligation. 5′ capped sense RNAs were synthesized using a construct encoding the transposase and the mMessage mMachine kit (Ambion). 20 pg of the Fli1ep:hPziezo1 DNA construct and 20 pg of the transposase sRNA were simultaneously injected into embryos at the one cell stage.

### Morpholino and CRISPR/Cas9 control experiments

The Pz1b MO1 and MO2 were designed to block the post-transcriptional splicing of the Pz1b. As with Pz1b MO1 (Suppl.fig.S3.A,B), we also confirmed that splicing had been disrupted by performing RT-PCR on Pz1b MO2 morphants (Suppl.fig.S3.I,J). To further ensure that the observed phenotype is due to specific knockdown of Pz1b, we co-injected suboptimal concentrations of both Pz1b MO1 and Pz1b MO2. Subsequent *in situ* hybridization with a *cmlc2a* probe reveals that injection of either morpholino at suboptimal concentrations alone does not affect heart development (Suppl.fig.S3.K-N). However, co-injection of both morpholinos at these concentrations results in the same defective cardiac looping phenotype observed at optimal concentrations of either morpholino (Suppl.fig.S3.O,P). Lastly, we also endeavoured to knockout (KO) Pz1b by employing a CRISPR/Cas9 strategy. To achieve this, we designed a gRNA which targeted exon 11 of Pz1b. In this manner we were able to generate a mutant zebrafish line harbouring a 13bp substitution at position 1435 to 1437 in Pz1b resulting in a frameshift and premature stop codon after 609 amino acids. This line was designated *piezo1b^mmr5^*. Unfortunately, homozygous *piezo1b^mmr5/ mmr5^* embryos did not display any discernible phenotype. A recent report has indicated that KO of the other *piezo1* zebrafish paralogue *piezo1a* also does not produce any discernible phenotype ^2^. Although it is possible that zebrafish do not require *piezo1* at all for development, based on the drastic consequences observed in mammals and humans when *piezo1* is KO or mutated, this seems unlikely ^3–5^. We reasoned that the most likely explanation for the lack of phenotype was due to compensation, as has been reported in other zebrafish KO lines where the expected phenotype was not observed^6^. To assess this possibility we followed the recently developed morpholino guidelines^7^ and injected the Pz1b morpholino into embryos from an incross of the *piezo1b^mmr5/mmr5^* line. If compensation has occurred, we should expect that the Pz1b morpholino would have little effect in homozygous *piezo1b^mmr5/ mmr5^* embryos. Indeed, we were able to observe an obvious oedema and defective cardiac looping in wildtype Pz1b morphant embryos (the embryos were produced by incrossing a *piezo1b* ^+/+^ line which was established in parallel to the KO line to maintain the same genetic background), however this defect was absent in *piezo1b^mmr5/mmr5^* morphants (Suppl.fig.S4.A,B). To analyse this difference in more detail, we performed a nitric oxide (NO) (DAF-FM-DA) assay to assess the production of this molecule in the OFT. Our analysis indicates that there is a significant decrease in the production of NO in the OFT in WT Pz1b morphant embryos compared to sibling WT controls (Adjusted P Value= <0.0001) (Suppl.fig.S4.C,D,G). However, there was no significant decrease in OFT NO production in *piezo1b^mmr5/ mmr5^* embryos injected with the Pz1b MO compared to sibling *piezo1b^mmr5/ mmr5^* controls (Adjusted P Value= 0.0212) (Suppl.fig.S4.E,F,G). This indicates that in *piezo1b^mmr5/ mmr5^* zebrafish embryos, Pz1b is compensated and thus the Pz1b morpholino has little effect. How it is compensated we are unable to tell at this juncture. This result also confirms that the phenotype we observe in Pz1b morphants is specific to knockdown of Pz1b and is not due to off target effects.

### Zebrafish cardiac performance analysis

The heart is at the centre of the circulatory system and consequently any reduction in its performance will have wide ranging consequences. Using the μZebraLab™ system from ViewPoint ^8^, we were able to determine that Pz1b morphant embryos had a significantly lower heart rate (160.4bpm +/−3.4SEM) than WT embryos (187.5bpm +/− 0.65SEM) (Suppl.fig.2.E). To investigate this phenotype further, we next analysed a number of blood flow parameters using the ViewPoint ZebraBlood™ system ^8^. In this manner we were able to determine that the mean blood flow in Pz1b morphants was significantly lower than that of un-injected controls (Suppl.fig.2.F). Further analysis of Pz1b morphants indicates that the maximal blood flow (corresponding to cardiac systole) is also markedly reduced when compared to the control embryos (Suppl.fig.2.G). Similarly, the average stroke volume is also significantly lower in the Pz1b morphants when compared to un-injected controls (Suppl.fig.2.H). The reduced blood flow rate associated with systole could also be an indicator that cardiac contractility is compromised in Pz1b morphants. To test this we made high-speed video recordings of either wildtype un-injected embryos or Pz1b morphants. Subsequent frame by frame analysis allowed us to measure the distance that the ventricular wall contracts during a cardiac cycle (Suppl.fig.6,B). In this manner, we found that ventricular contraction in Pz1b morphants was increased resulting in a reduced end systolic volume (ESV) (Suppl.fig.2.I). Furthermore, we were also able to observe blood regurgitation occurring between the atrium and ventricle at the end of systole (movies S1 and S2). Taken together, the decreased ESV and cardiac output observed in Pz1b morphants suggests the presence of OFT stenosis.

### Zebrafish strains and husbandry

Zebrafish were maintained under standardized conditions and experiments were conducted in accordance with local approval (APAFIS#4054-2016021116464098 v5) and the European Communities council directive 2010/63/EU. Embryos were staged as described ^9^. The *Tg(fli1a:GFP)y1Tg* was provided by the CMR[B] *Centro de Medicina Regenerativa de Barcelona*. The double transgenic line *Tg(fli1a:GFP)y1;Tg(cmlc2a:RFP*) was generated in house. All larvae were euthanised by administration of excess anaesthetic (Tricaine).

### Morpholinos and injections

Morpholino oligonucleotides (MOs) were obtained from Gene Tools (Philomath, OR, USA) and injected into one-cell stage embryos. The Pz1b MO1 and MO2 target the intron 7/exon 8 and the last exon/intron splice site of *piezo1b* (GenBank: KT428876.1) respectively (supplemental information).

### Embryonic zebrafish endothelial primary cell culture

3dpf *Tg(fli1a:GFP)y1* embryos were placed in a microtube and pipetted up and down 10 times to remove the yolk sac. Digestion was performed in digestion buffer (2.5mg/mL trypsin, 1mM EDTA, 0.16mg/mL tricaine in PBS) in the incubator (28°C) and monitored every 10 minutes and pipetted up and down to dissociate cells mechanically. The reaction was stopped by adding a same volume of 10% FBS, 2mM CaCl_2_. Cells were pelleted by centrifugation 3 min 3000rpm, washed with culture medium (L-15, 5% FBS, 1% penicillin/streptavidin) and re-suspended in culture medium. Cells were plated in a 35mm Petri dish (5 embryos/2mL/per dish) and placed in an atmosphere of 95% air/5% CO_2_ at 30 °C for a minimum 2 hours prior to patch clamp experiments.

### Cell surface quantification

For membrane expression quantification of the WT and variant PIEZO1, fluorescence intensity (grey value) was measured across a line from the outside to the inside of the cell for GFP and RFP using ImageJ.

### PIEZO1 3D structure

Residues K2528 (Magenta) and Y2038 (Green) identified in this study as subject to genetic variability are represented as spheres. Images were produced using PyMOL ™ Molecular Graphics System, Version 1.8.6.0.

### CRISPR/Cas9

Pz1b target sequences were identified using ZiFiT online software ^10^. The Pz1b gRNA was synthesised from a DNA string template from Invitrogen using the Megashortscript T7 kit (Ambion) and purified by phenol/chloroform extraction. 150pg of Pz1b gRNA was co-injected with nls-Cas9 protein (N.E.B). Embryos were bred to adulthood and initially screened for mutations in Pz1b using a previously described T7 assay ^11^. The founder of the *piezo1b^mmr5^* mutant line was verified by Sanger sequencing and outcrossed to AB wildtype. F1 *piezo1b*^*mmr5*/+^ adults were identified using the T7 assay and verified by Sanger sequencing.

### BDM, PHZ and GsMTx4 treatment

BDM (10 or 15mM, Sigma-B0733), PHZ (0.1mg/mL, Sigma-114715) and GsMTx4 (1μM, Smartox Biotechnology) were added to the embryo medium 2 hours, 2 days and 4 hours before analysis respectively.

### TRIM, PKI166 and AG478 treatments

Fir the ISH analysis the NO inhibitor TRIM (0.25 mM, Sigma-T7313) was added to the embryo medium at 48hpf. PKI166 AND AG1478 treatments were performed as described ^12^. For the aortic valves analysis, TRIM (0,01mM) was added to the embryo medium at 24hpf and refreshed daily until 7dpf.

### Electrophysiology

Each current was evaluated using an Axiopatch 200B amplifier (Axon Instrument, USA), low-pass filtered at 3kHz and digitized at 2kHz using a 12-bit analog-to-digital digidata converter (1440A series, Axon CNS, Molecular devices, USA). Results are expressed as mean ± standard error of the mean (SEM). Patch clamp pipettes were pulled using vertical puller (PC-10, Narishige) from borosilicate glass capillaries and had a resistance 1.2-2MΩ for the inside out currents. For the whole cell attached currents, the bath solution contained (in mM) 155 KCl, 3 MgCl_2_, 5 EGTA and 10 HEPES adjusted to pH7.2 with KOH. The pipette solution contained (in mM) 150 NaCl, 5 KCl, 2 CaCl_2_, and 10 HEPES adjusted to pH7.4 with NaOH.

### *In situ* hybridisation

The *Pz1b* probe was generated by cloning a 2kb fragment from a 3dpf zebrafish cDNA library into pGEMT. The *cmlc2a* probe was generated from pBSK *cmlc2a* containing the full length *cmlc2a* gene. The *elnb* probe was generated by cloning a 1kb fragment from a 3dpf zebrafish cDNA library into pGEMT. The *acana* probe was generated as described previously. ISH on larvae were performed as described previously^13^. Anti-sense probes were synthesized as described previously ^14^. High-resolution fluorescent double whole-mount *in situ* hybridization were performed as described previously ^15^.

### DAF labelling

To reveal the presence of NO, embryos were incubated in E3 medium containing 5μM DAF-FM DA (Life Technologies, D23842), for 1 hour in the dark at 28°C. Fluorescence intensities of the OFT were measured using ImageJ software.

### Immunohistochemistry

Transfected cells were fixed in 4% PFA for 20min and blocked with PBS, 2mg/mL BSA, 2% lamb serum for 45min at room temperature. Cells were then incubated in the same solution with primary rabbit anti-Piezo1 antibody (Proteintech 15939-1-AP, 1/100) for 3h followed by donkey Alexa Fluor 594-anti rabbit (Jackson ImmunoResearch Laboratories, 1/200) 2h at room temperature.

### RT-PCR analysis

30hpf embryos non-injected or injected with 5ng of *piezo1b* MO1 or with 5 or 10ng of *piezo1b* MO2 were collected and RNA was isolated with TRIzol^®^ Reagent. RT-PCR was performed on 1μg of RNA using oligo-dT primers and amplification PCR was performed using *piezo1b* specific primers

The sequences of the Pz1b specific primers are the following:

For (MO1): 5′-GTTTTGATCGTAACGTCAAA-3′
Rev (MO1): 5′-TCAGTTGTTTGTGTCTCTTT-3′
For (MO2): 5′-ACGGAGTCTATAAGAGCATC-3′
Rev (MO2): 5′-TCATTTGTTTTTCTCTCGCG-3′

### Cardiovascular parameters analysis

To determine the cardiovascular parameters, we utilized TZebraLab™ software from ViewPoint which has been developed specifically for this purpose. All experiments were performed as described previously ^8^. To determine the mean blood flow, the average of the blood flow values was calculated from a 10 second time frame (1,300 frames). For the maximum and minimum blood flow, averages of highest and lowest peak values were calculated during 20 heartbeats. Stroke volume was calculated by dividing the average blood flow (nL/sec) by the heart beat per second (BPM/60). To assess the heart contractility, movies were recorded at 120fps and measurements were made using ImageJ software.

### Confocal Imaging

Still images of fixed or anesthetized embryos were recorded using a ProgRes CF colour camera (Jenoptik) mounted on a SZX16 stereomicroscope (Olympus). Fluorescent embryonic heart images were acquired with a Leica TCS SP8 inverted confocal laser scanning microscope with a 20X or 40X oil immersion objectives. Maximal projections and analysis were performed with Imaris or ImageJ softwares. OFT movements were recorded using a high speed 12kHz resonant scanner. Image acquisition and image analysis were performed on workstations of the Montpellier RIO Imaging facility of the Arnaud de Villeneuve/IFR3 campus.

### PCGC data analysis

Whole exomic data for individual probands was downloaded from the NIH/dbGaP repository (The Pediatric Cardiac Genomics Consortium (PCGC)). The data was subsequently converted into the Fastq format using the SRA toolkit provided by NIH/dbGaP. The exomic data was aligned to the human reference genome and SNPs associated with *PIEZO1* were identified using the Integrative Genomics Viewer ^16^. Patients’ data analysis was approved by the INSERM Institutional Review Board (IRB00003888 – Opinion number 15-221).

## Supplemental figure legends

**Figure S1.**
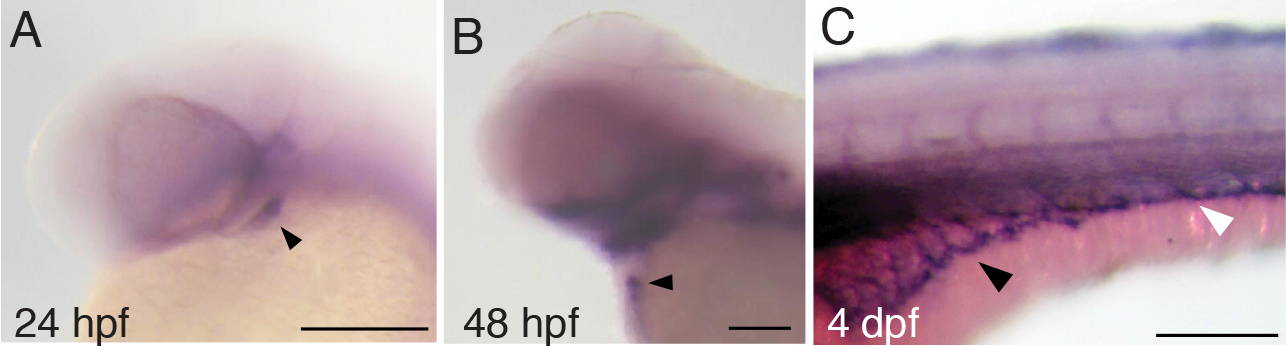
Pz1b is expressed in endothelial cells. (**A-C**) ISH using an antisense Pz1b probe. (**A,B**) Pz1b expression can be detected in the heart of zebrafish embryos (black arrowhead) at 24hpf (**A**), 48hpf (**B**). (**C**) Pz1b is also expressed in the developing vasculature present on the yolksac (black arrowhead) and in the tail (white arrowhead) of 4dpf zebrafish embryos. Scale bars: (**A-C**) 100μm

**Figure S2.**
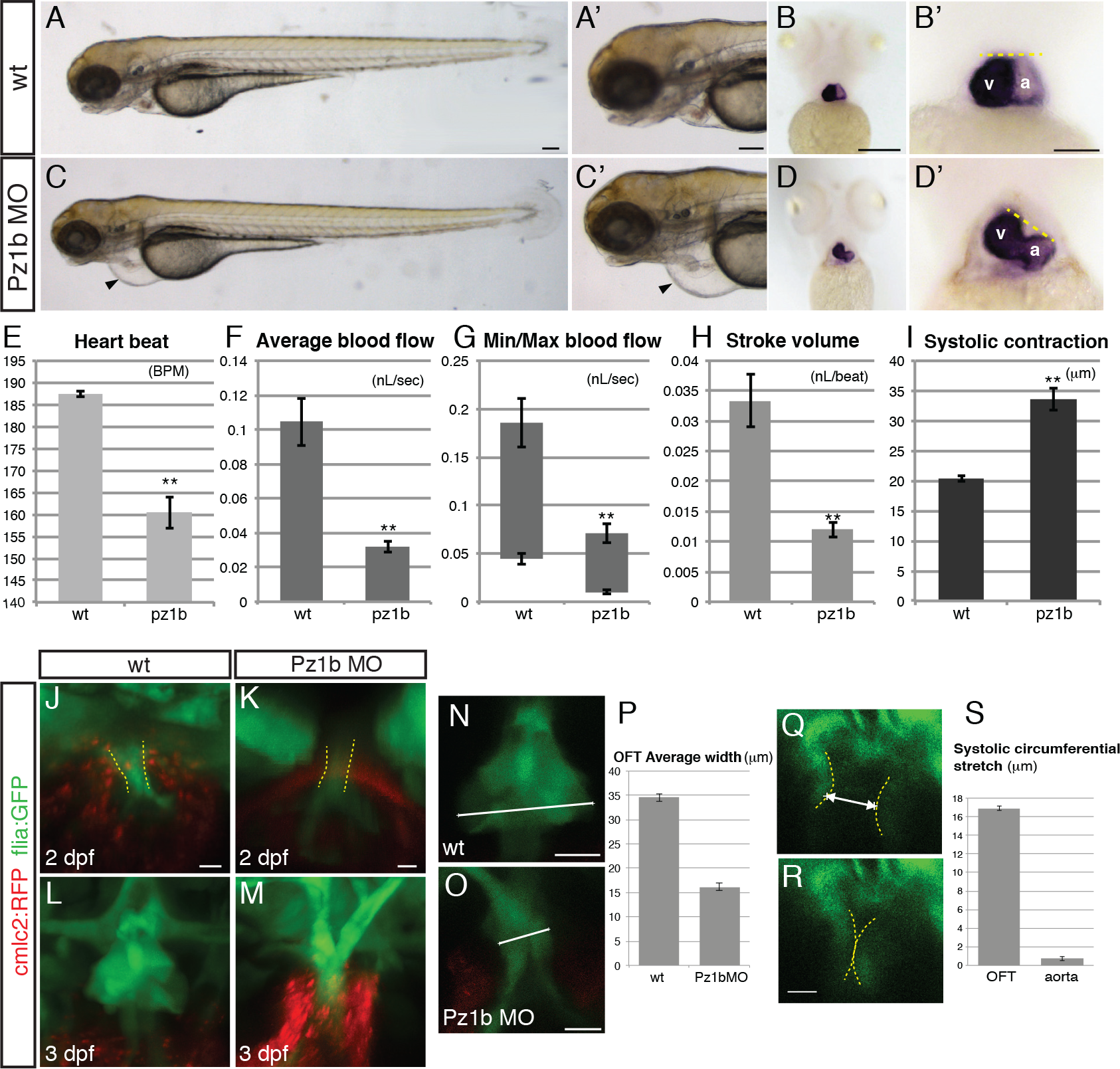
*Piezo1b* is involved in cardiovascular development. (**A**) Un-injected 72hpf zebrafish embryo, (**A’**) the same embryo at higher magnification. (**C**) Pz1b MO 72hpf morphant displaying pericardial oedema and defective cardiac looping (black arrowhead), (**C’**) the same embryo at higher magnification. (n=48/50). (**B,B’,D,D’**) ISH using an antisense *cmlc2a* probe on (**B**) 72hpf un-injected zebrafish embryos, (**B’**) the same embryo at higher magnification, the yellow dashed line above the ventricle (v) and atrium (a) is an indicator for the degree of cardiac looping. (**D**) 72hpf Pz1b MO morphant embryo, (**D’**) the same embryo at higher magnification, note the yellow dashed line is no longer horizontal indicating a defect in cardiac looping (n=48/50). (**E**) Graph depicting the average heart rate in beats per minute (BPM) of WT (n=8) and Pz1b morphants (n=8). (**F**) Graph depicting the average blood flow rate in nano litres per second (nL/sec) of WT (n=8) and Pz1b morphants (n=8). (**G**) Graph depicting the average minimum/maximum blood flow rate in nano litres per second (nL/sec) of WT (n=8) and Pz1b morphants (n=8). (**H**) Graph depicting the average stroke volume in nano litres per heart beat (nL/beat) of WT (n=8) and Pz1b morphants (n=8). (**I**) Graph depicting the average ventricular contractile distance in micro metres (μM) of WT (n=5) and Pz1b morphants (n=5). (**J-M**) Maximal projections of 48hpf (**J,K**) and 72hpf (**L-M**) *Tg(fli1a:GFP)y1;Tg(cmlc2:RFP*) OFT from WT (**J,L**) and Pz1b morphants (**K,M**). Yellow dashed lines delineate the OFT position at 48hpf (**J,K**). (**N,O**) Maximal projections of a 72hpf *Tg(fli1a:GFP)y1* OFT from WT (**N**) and Pz1b morphants (**O**). The white line indicates the width of the OFT immediately adjacent to the myocardium. (**P**) Graph showing the average size of the OFT of WT and Pz1b MO1 injected embryos (n wt=7, n Pz1bMO1=11). (**Q,R**) Resonance laser z-stack imaging of a 48hpf *Tg(fli1a:GFP)y1* OFT at 250ms interval, the yellow dashed lines delineate the inside of the OFT. The white double headed arrow indicates the OFT inside diameter during systole (**Q**). (**S**) Graph showing the difference between OFT and aorta inside diameters between diastole and systole (n=4; 3 contractions for each). All graphs (* p<0,05, ** p<0.01, student’s t-test),Scale bars (**A-B’**) 100μm (**J-O** and **Q,R**) 10μm.

**Figure S3.**
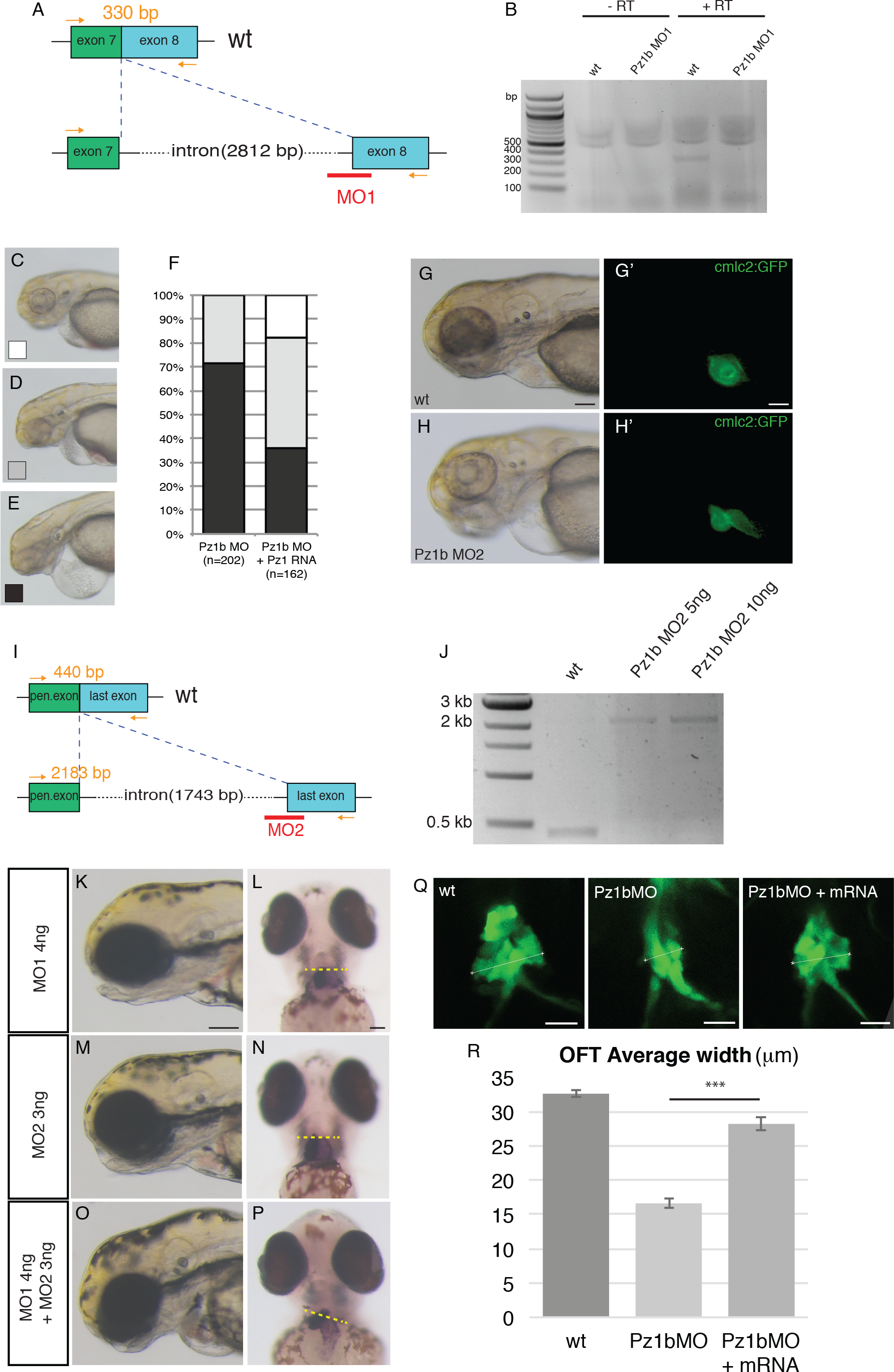
Pz1b experimental controls. (**A,B**) RT-PCR analysis of Pz1b MO1 splice morphant embryos. The Pz1b MO1 targets the intron 7/exon 8 splice site. Primers designed against exons 7 and 8 (arrows in (**A**)) are able to amplify the correct 330bp fragment (lower band in **B**) from WT embryo cDNA (lane 3). With morphant embryo cDNA, the final intron has not been spliced out (lane 4) indicating *piezo1b* has been disrupted. The first two lanes correspond to the negative control without reverse transcriptase (RT). (**C-F**) Mouse *Piezo1* RNA can rescue the Pz1b morphant phenotype. Representative images of the severity of cardiac oedema: (**C**) no oedema, (**D**) moderate oedema, (**E**) severe oedema. (**F**) Graph indicating the percentage of embryos displaying either no oedema (white), moderate (grey) or severe (black) in either Pz1b morphant embryos (Pz1b MO) or Pz1b morphants co-injected with 20pg of mPz1 RNA (Pz1b MO+Pz1 RNA) n= the cumulative number of embryos from 5 separate experiments. (**G**) Brightfield image of an un-injected 3dpf *Tg(cmlc2a:GFP*) zebrafish embryo, (**G’**) fluorescent image of the same embryo, GFP can be detected in the heart (green). (**H**) Brightfield image of an Pz1b MO2 injected 3dpf *Tg(cmlc2a:GFP*) zebrafish embryo, (**H’**) Fluorescent image of the same embryo, GFP can be detected in the heart (green), note the linear appearance of the chambers (n=38/40). (**I,J**) RT-PCR analysis of Pz1b MO2 morphant embryos. The Pz1b MO2 targets the last intron/exon splice site (**I**). Primers designed against the final two exons (arrows in (**I**)) are able to amplify the correct 440bp fragment (lower band) from wildtype embryo cDNA (lane 1). (**J**) With morphant embryo cDNA, the final intron has not been spliced out (upper 2,183bp band in lanes 2 and 3) indicating *piezo1b* has been disrupted. (**K**) Brightfield image of a 3dpf embryo injected with suboptimal (4ng) concentration of Pz1b MO1. (**L**) *In situ* hybridization using an antisense *cmlc2a* probe indicates heart looping is normal (n=20/20). (**M**) Brightfield image of a 3dpf embryo injected with suboptimal (3ng) concentration of Pz1b MO2. (**N**) *In situ* hybridization using an antisense *cmlc2a* probe indicates heart looping is normal (n=20/20). (**O**) Brightfield image of a 3dpf embryo injected with suboptimal concentration of Pz1b MO1 (4ng) and Pz1b MO2 (3ng). (**P**) *In situ* hybridization using an antisense *cmlc2a* probe indicates heart looping is disrupted (n=20/22). **(Q-R**) Mouse *Piezo1* RNA can rescue the Pz1b morphant OFT phenotype. Representative images of the OFT in WT larvae (wt), Pz1b morphants (Pz1b MO) and Pz1b morphants co-injected with mPz1 RNA (Pz1bMO+mRNA). The white line indicates where the width of the OFT was measured. (**R**) Graph indicating the average width of the OFT in WT (n=10), Pz1b morphants (n=10) and Pz1b morphants co-injected with mPiezo1 mRNA (n=12) (* p<0,05, ** p<0.01, student’s t-test). Scale bars (**G,G’,K,L**) 100μm.

**Figure S4.**
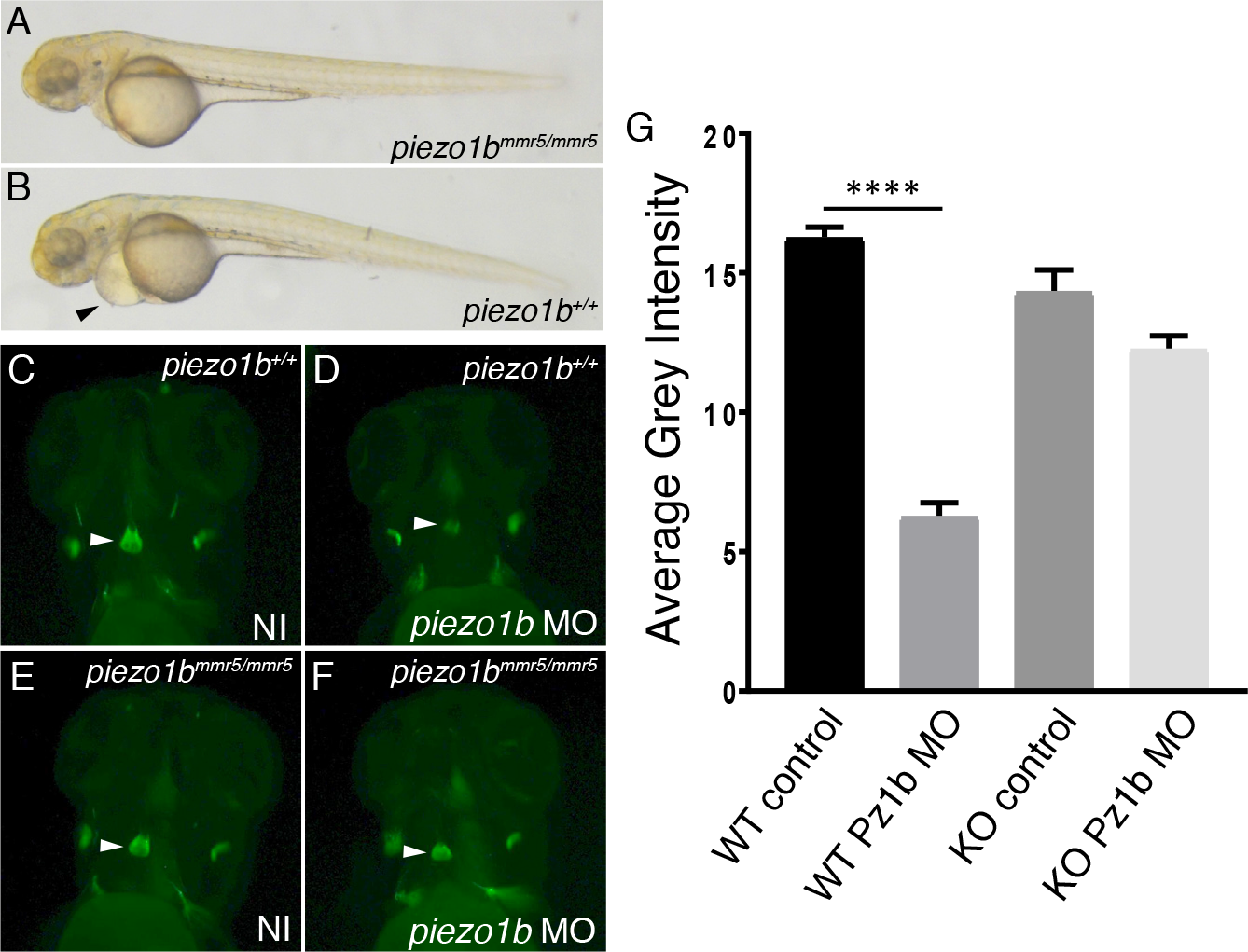
Homozygous *piezo1bmmr5/ mmr5* knockout zebrafish larvae are compensated. (**A**) Brightfield image of a 3dpf *piezo1b^mmr5/ mmr5^* zebrafish embryo injected with Pz1b MO. (**B**) Brightfield image of a 3dpf *piezo1b^+/+^* zebrafish embryo injected with Pz1b MO (black arrow head indicates cardiac oedema). (**C**) A 3dpf *piezo1b^+/+^* zebrafish embryo treated with DAF-FM DA to reveal NO production in the OFT (white arrow head indicates the OFT). (**D**) A 3dpf *piezo1b^+/+^* zebrafish embryo injected with Pz1b MO and treated with DAF-FM DA to reveal NO production in the OFT (white arrow head indicates the OFT). (**E**) A 3dpf *piezo1b^mmr5/ mmr5^* zebrafish embryo treated with DAF-FM DA to reveal NO production in the OFT (white arrow head indicates the OFT). (**F**) A 3dpf *piezo1b^mmr5/ mmr5^* zebrafish embryo injected with Pz1b MO and treated with DAF-FM DA to reveal NO production in the OFT (white arrow head indicates the OFT). (**G**) Graph showing mean grey intensity values (WT control n=48, WT Pz1b MO n=51, KO control n=35, KO Pz1b MO n=39). Error bars indicate SEM, ANOVA and Sidak’s multiple comparisons test ****P<0.0001.

**Figure S5.**
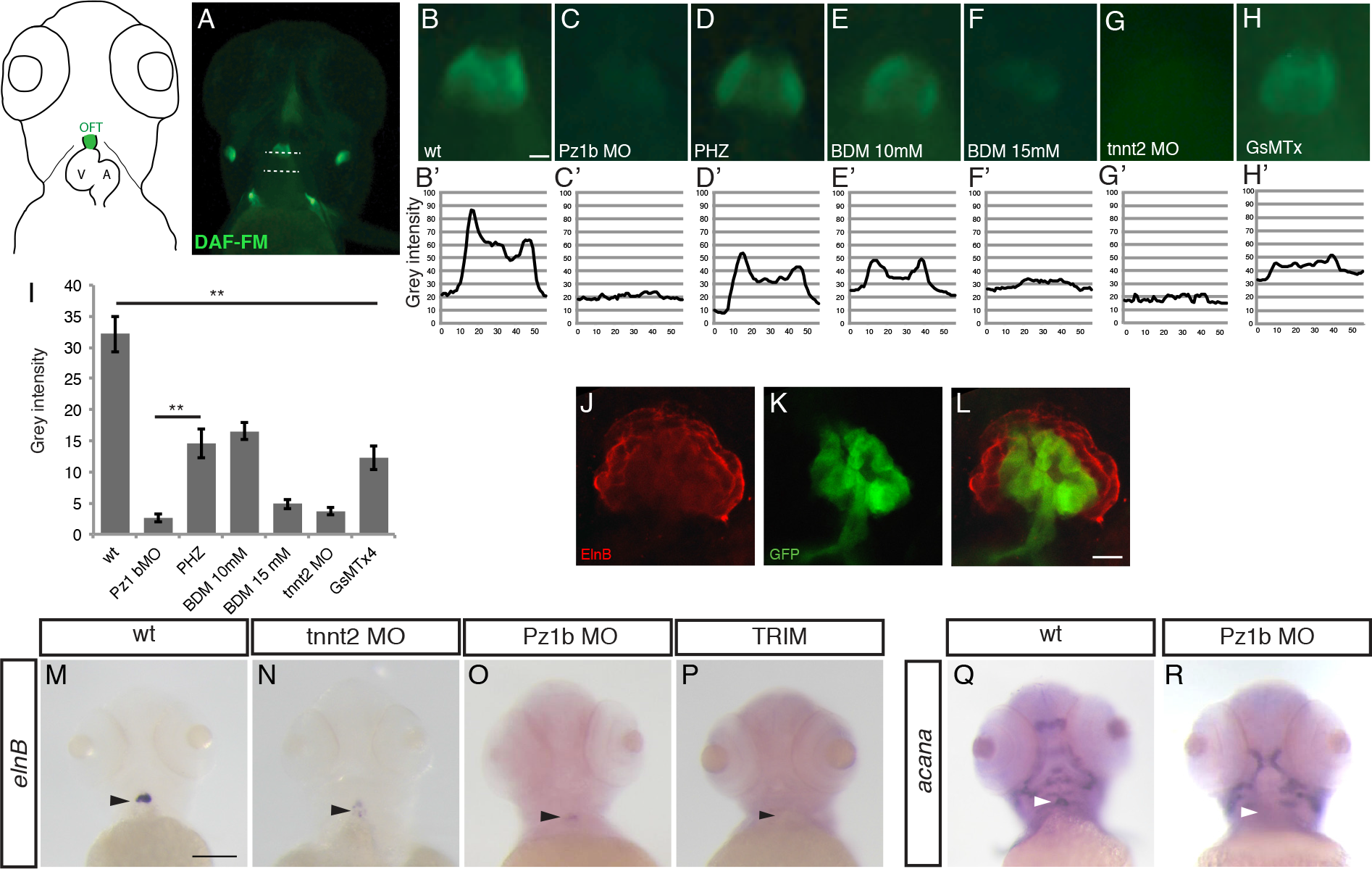
*Piezo1b* regulates nitric oxide production and extracellular matrix composition in the outflow tract. (**A**) WT 72hpf zebrafish embryo treated with DAF-FM DA to reveal NO production in the OFT (bisected by upper dashed white line). Dashed white lines indicate the regions where grey scale intensity was measured, the upper line bisects the OFT and the lower line is used to measure the background intensity. (**B**) Higher magnification image of the OFT of an un-injected embryo assayed with DAF-FM DA, (**B’**) graph showing the grey scale intensity across the OFT. (**C-G’**) OFT images and grey scale graphs of a 72hpf Pz1b morphant (**C,C’**), PHZ treated embryo (**D,D’**), 10mM BDM treated embryo (**E,E’**), 15mM BDM treated embryo (**F,F’**), tnnt2 morphant (**G,G’**) or a GsMTx4 treated embryo (**H,H’**). (**I**) Graph showing the average grey scale intensity of un-injected controls (wt), Pz1b morphants (Pz1b MO), PHZ treated embryos (PHZ), embryos treated with 10mM BDM (BDM 10mM), embryos treated with 15mM BDM (BDM 15mM), tnnt2 morphant or GsMTx4 treated embryos (GsMTx4) (n=10 for each condition). **(J-L)** Single z-stack confocal images of the OFT following an immunostaining using an anti-ElnB antibody (**J**, red) on 3 dpf *Tg(fli1a:GFP)y1* embryos (**H**, green). The merged image (**L**) indicates that the cells producing ElnB are not endothelial which produce GFP. **(M-P)** *In situ* hybridization against *elnB* in 3 dpf WT, *tnnt2*, Pz1b morphants and TRIM treated larvae. **(Q,R)** *In situ* hybridization against *acana* in 3 dpf WT, and Pz1b morphants. Scale bars (**A**) and **(M-R)** 100μm; (**B**) 20μm; **(J-L)** 10μm. All graphs show mean values. Error bars indicate SEM, ANOVA and Dunnet’s multiple comparisons test **P<0.01(upper line, all samples), student’s unpaired homoscedastic two tailed t-test **P<0.01 (lower line Pz1b vs PHZ).

**Figure S6.**
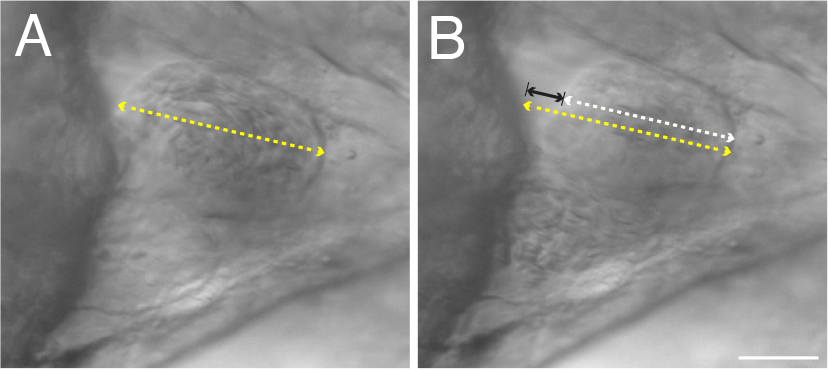
Pz1b knockdown perturbs systolic contraction. (**A,B**) Representative images of contraction analysis using ImageJ software. The distance between the two ventricular walls is first measured at the end of diastole (yellow double headed arrow)(**A**), (**B**) the distance is again measured at the end of systole (white double headed arrow, the difference is indicated by the black double headed arrow).

**Figure S7.**
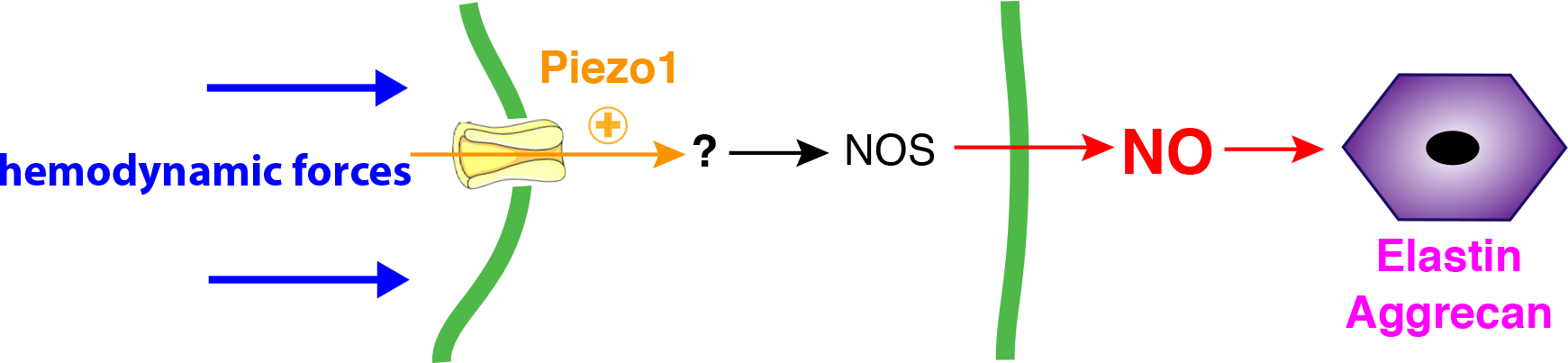
Model of Pz1b signalling pathway. A cartoon schematic depicting the authors proposed signalling cascade. Hemodynamic forces in the OFT cause Piezo1 to open releasing cation into the endothelial cells cytoplasm. This induces an as of yet unidentified process to activate NOS resulting in the release of NO. This in turn induces the cells which ensheathe the OFT to begin producing ECM components such as Elastin and Aggrecan.

**Movie S1. The beating heart of a 3dpf wildtype embryo.**

High speed video recording capturing the beating heart of a 3dpf wildtype embryo. 5 heart beats are shown at 120fps then the same 5 heart beats are shown at 40fps. Blood can be seen entering the atrium and passing to the ventricle before ejection.

**Movie S2. The beating heart of a 3dpf Pz1b morphant.**

High speed video recording capturing the beating heart of a Pz1b morphant. 5 heart beats are shown at 120fps then the same 5 heart beats are shown at 40fps. Following the contraction of the ventricle blood can be seen regurgitating back into the atrium.

**Movie S3. OFT dynamism of a 48hpf *Tg(fli1a:GFP)y1* embryo.**

Resonance laser imaging capturing the dynamic movement of the OFT during systole and diastole. Frames were acquired every 50ms.

**Movie S4. Dorsal aorta dynamism of a 48hpf *Tg(fli1a:GFP)y1* embryo**.

Resonance laser imaging capturing the dynamic movement of the dorsal aorta during systole and diastole. Frames were acquired every 130ms.

**Movie S5. Aortic valves in a 7dpf wildtype zebrafish larvae.**

2 photon imaging of a BODIPY labelled 7dpf zebrafish larvae reveals the valves dynamically moving during the cardiac cycle.

**Movie S6. Aortic valves in a 7dpf Pz1b morphant zebrafish larvae.**

2 photon imaging of a BODIPY labelled 7dpf Pz1b morphant zebrafish larvae reveals the valves dynamically moving during the cardiac cycle.

**Movie S7. Aortic valves in a 7dpf *notch1b* mutant zebrafish larvae**

2 photon imaging of a BODIPY labelled 7dpf *notch1b* mutant zebrafish larvae reveals the valves dynamically moving during the cardiac cycle.

**Movie S8. Echocardiogram of the aortic valve from the Y2022H proband.**

**Movie S9. Echocardiogram of the aortic valve from the K2502R proband.**

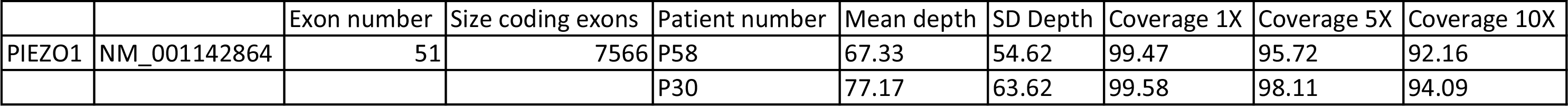

| Coverage 20X | Coverage 30X |
| --- | --- |
| 82.9 | 72.55 |
| 85.8 | 75.95 |

